# Electroencephalographic Signatures of Canine Cognitive Dysfunction

**DOI:** 10.1101/2022.01.28.478033

**Authors:** Alejandra Mondino, Mary Gutiérrez, Camila González, Diego Mateos, Pablo Torterolo, Natasha Olby, Luis Delucchi

## Abstract

Canine cognitive dysfunction (CCD) is a highly prevalent neurodegenerative disease considered the canine analog of Alzheimer’s disease (AD). Unfortunately, CCD cannot be cured. However, early therapeutic interventions can slow the progression of cognitive decline and improve quality of life of the patients; therefore, early diagnosis is ideal. In humans, electroencephalogram (EEG) findings specific to AD have been described, and some of them have successfully detect early stages of the disease. In this study we characterized the EEG correlates of CCD, and we compared them with the EEGs of healthy aging dogs and dogs at risk of developing CCD. EEG recordings were performed in 25 senior dogs during wakefulness. Dogs were categorized in normal, at risk of CCD or with CCD according to their score in the Rofina questionnaire. We demonstrated that, quantitative EEG can detect differences between normal dogs and dogs with CCD. Dogs with CCD experience a reduction in beta and gamma interhemispheric coherence, and higher Joint Lempel Ziv complexity. Dogs at risk of developing CCD, had higher alpha power and interhemispheric coherence, making these features potential markers of early stages of the disease. These results demonstrate that EEG could be an additional biomarker that can contribute to the diagnosis of CCD, and reinforce the CCD as a translational model of AD.

## Introduction

Canine cognitive dysfunction (CCD) is a highly prevalent neurodegenerative disease that affects older dogs and is characterized by progressive signs of altered mentation, deficits in memory and learning, changes in socialization and in the sleep-wakefulness cycle (Dewey et al., 2019; Rofina et al., 2006). Due to its clinical and histopathological similarities, CCD is considered the canine analog of Alzheimer’s disease (AD) (Prpar Mihevc and Majdic, 2019; Schutt et al., 2016). Unfortunately, at this time, it is not possible to cure CCD, but certain therapies can slow the progression of cognitive decline and improve quality of life of the patients (Campbell et al., 2001; Pero et al., 2019; Pop et al., 2010; Zakosek Pipan et al., 2021). An early diagnosis is ideal because therapeutic impact is greater if initiated early in disease course (Landsberg et al., 2012; Osella et al., 2007). Owner-based questionnaires are the most employed diagnostic tools; though, they represent a subjective and indirect evaluation of the disease and their diagnostic sensitivity and specificity has not been evaluated against the gold standard for diagnosis, histopathology (Madari et al., 2015; Rofina et al., 2006; Salvin et al., 2011; Schütt et al., 2015). Additionally, magnetic resonance imaging (MRI) of the brain can show abnormalities suggestive of CCD such as brain atrophy, leukoaraiosis and a reduction of the interthalamic adhesion thickness (Hasegawa et al., 2005; Scarpante et al., 2017). However, due to costs and potential risk of anesthesia, owners are sometimes reluctant to have an MRI performed in their aging dogs. In the past few years, research has focused on the development of additional tools that could detect and quantify early stages of the disease, such as plasma and cerebrospinal fluid biomarkers (Gonzalez-Martinez et al., 2011; Panek et al., 2020; Panek et al., 2021; Rosado et al., 2012; Smith et al., 2021; Van Bourg et al., 2021) or cognitive tests (Chapagain et al., 2017; Heckler et al., 2014; Hoel et al., 2021; Mongillo et al., 2013; Piotti et al., 2018). Nevertheless, biomarkers analyses are not yet easily available, and are not disease specific. In humans, several correlates of the electroencephalogram (EEG) have been associated with AD, and some of them have successfully detect early stages of the disease and predict progression to AD in mild cognitive impaired patients (Jeong, 2004; Kowalski et al., 2001; Poil et al., 2013). EEG oscillations contain complex frequency spectra that can be analyzed by power spectrum analysis. The power of each frequency reflects the degree of local synchronization of the extracellular potential at that specific frequency (Buzsaki et al., 2013). Additionally, long-range synchronization at each frequency between different cortical areas can be measured by means of coherence analysis (Castro et al., 2014; Torterolo et al., 2019). Power and coherence highly change between different behavioral states such as, sleep and wakefulness, altered states of consciousness and during unconsciousness (Gonzalez et al., 2020; Mondino et al., 2020; Mondino et al., 2021a; Mondino et al., 2022). High frequency oscillations such as beta and gamma are considered to have an important role in cognition and memory (Mably and Colgin, 2018; Piai et al., 2015). In AD patients, high frequency oscillations show a decrease in power spectrum and connectivity between cortical areas, while power of slow oscillations increases in patients with AD (Dauwan et al., 2016; Jeong, 2004; Kowalski et al., 2001; McBride et al., 2015). In addition to this, is it possible to evaluate the complexity of the EEG signal by different analysis such as Lempel Ziv complexity (Lempel and Ziv, 1976). Furthermore, while some contradictory results have been found, most studies show a decrease in EEG complexity (Abasolo et al., 2007; Liu et al., 2016; Mizuno et al., 2010; Simons and Abásolo, 2017; Sun et al., 2020). The EEG changes associated with CCD have not been evaluated yet and they could provide novel non-invasive biomarkers for the early detection and quantification of the disease. The main goal of this study was to characterize the EEG correlates of CCD and to compare them with healthy aging dogs and dogs at risk of developing CCD.

## Materials and methods

### Dogs

Family-owned dogs, older than 8 years old were recruited for this study through the Universidad de la República College of Veterinary Medicine, Montevideo, Uruguay. All dogs had a physical and neurological examination and Complete Blood Count (CBC) and serum chemistry was performed. Dogs were excluded if they had a history of epileptic seizures, showed signs of concurrent neurological disease (other than behavioral changes compatible with CCD) or CBC and chemistry results indicative of systemic disease. All the experimental procedures were approved by the Institutional Animal Care Committee (Protocol number 111900-000702-21).

### Owner-based questionnaire

Owners were asked to complete the Rofina questionnaire (Rofina et al., 2006). From the different canine dementia questionnaires available, this is the only one that has been validated in Spanish language (Gonzalez-Martinez et al., 2011). The questionnaire involves questions in 10 domains: appetite, drinking, elimination-behavior, sleep-wakefulness cycle, aimless behavior, interaction, perception, disorientation, memory, and personality changes. A Rofina dysfunction Score (RDS) is obtained by summing the score in each question, with higher values indicating higher levels of cognitive dysfunction (Rofina et al., 2006). Based on their RDS, dogs were categorized into three groups, Normal (10-14), At risk of developing CCD (15-21) or with CCD (>22) (Schütt et al., 2015).

### Electroencephalographic recordings

EEG recordings were performed using subcutaneous needle electrodes. Localization of the electrodes is schematized in Figure 1A. Active electrodes were placed at the left and right frontopolar (Fp1, and Fp2), frontal (F3, F4), parietal (P3, P4) and occipital cortex (O1, O2) (Pellegrino, 2004). These electrodes were referenced to the average signal of the right and left mastoid (A1, A2). A ground electrode was positioned at Fz (midline frontal). The signal was collected, pre-filtered and amplified, and digitalized with a sampling rate of 256 Hz using a AKONIC BIO-PC System (AKONIC, TM-Buenos Aires, Argentina) using the following settings: sensitivity 70 μV/cm, time constant 0.3 s, bandpass filter: 0.5 – 100 Hz, notch filter at 50 Hz. The impedance of all electrodes was always kept below 10kΩ.

**Figure 1.**
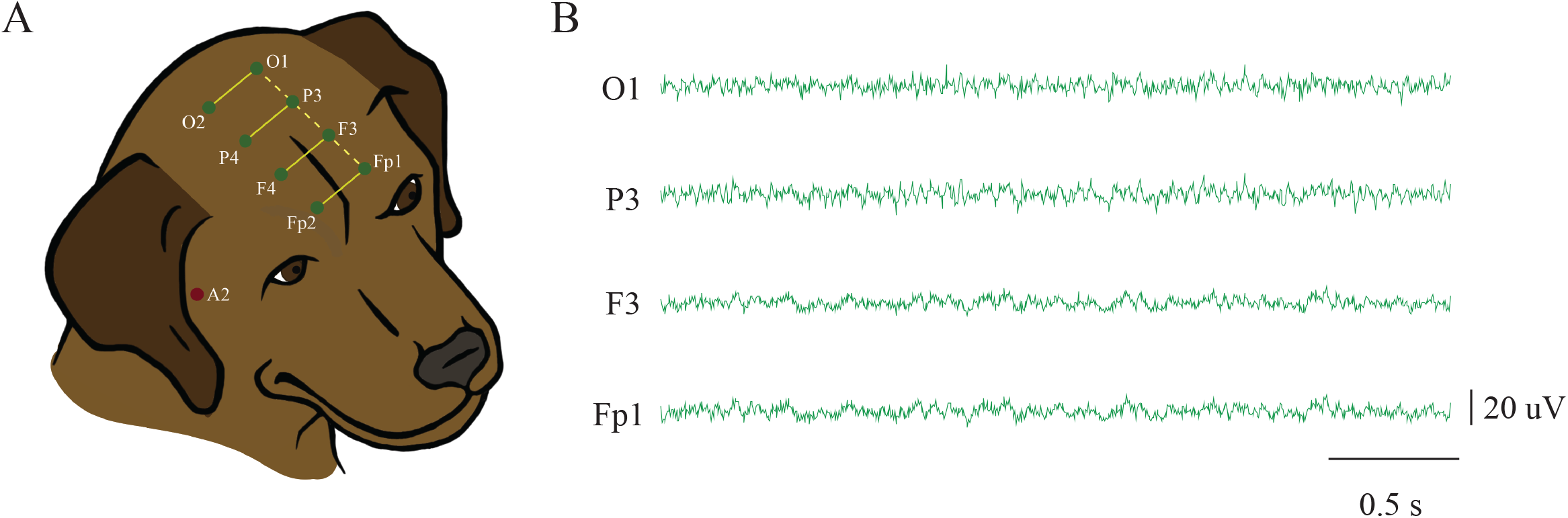
Electrode location for EEG recordings. **A.** The location of active electrodes is represented by green dots. We recorded the EEG activity of the frontopolar (Fp), frontal (F), parietal (P) and occipital (O) cortices. Odd numbers represent left cortices and even numbers represent right cortices. Active electrodes were referenced to the average signal of the right and left mastoid (A1, A2). A2 is represented by a red dot in the figure. Yellow lines show the combination of adjacent electrodes used to calculate interhemispheric (solid line) and intrahemispheric (dashed line) coherence. **B**. Representative EEG recordings of the left cortices in a dog with CCD.

### Data analysis

The EEG was recorded for at least seven minutes in waking animals. Every 5 s of recording were analyzed using Spike 2 software version 9.04 (Cambridge Electronic Design, Cambridge, UK) and the maximum number of artifact-free epochs where the dog was awake were selected for further analysis. Power spectrum was calculated using the *pwelch* built-in function in MATLAB version R2020a (The MathWorks Inc, Natick, MA, USA) with the following parameters: window = 1s, noverlap = 0, nfft = 512, fs = 256, which corresponds to employing 1s sliding windows with no window overlap and a 0.5 Hz resolution. Power values were calculated for delta (1 – 4 Hz), theta (4.5 – 8 Hz), alpha (8.5 – 12 Hz), sigma (12.5 – 16 Hz), beta (16.5 – 30 Hz) and gamma (30.5 – 45 Hz)(Kis et al., 2017). Relative power was calculated for each dog as the mean power of each frequency band over the total power (i.e., the sum of the power of all bands)(Mondino et al., 2021b).

Interhemispheric and intrahemispheric cortical coherence between adjacent channels (Figure 1A) were quantified with the *mscohere* MATLAB function with the same parameters and for the same frequency bands described above for the spectral power analysis. Coherence values were normalized by the Fisher’s z-transform(Miranda de Sá et al., 2009). Finally, we estimated the temporal complexity of the brain for the whole spectrum of frequencies (1 – 45 Hz) by means of Lempel-Ziv Complexity (LZC)(Lempel and Ziv, 1976). This measure is based on Kolmogorov complexity and calculates the minimal “information” contained in a sequence(Cover and Thomas, 2006; Schartner et al., 2015). To analyze LZC continuous sequences, such as the EEG signal, it needs to be discretized; in this work we employed the binarization by mean-value (Zozor et al., 2005). We also calculated the spatio-temporal complexity of the neurophysiologic activity, referred as joint Lempel-Ziv complexity (JLZC) (Zozor et al., 2005). For further details on the analysis please refer to Mondino et al. 2021 (Mondino et al., 2021a). Differences among groups in power and coherence for each frequency band, as well as for LZC and JLZC, were evaluated by means of one-way ANOVA followed by a Tukey *post-hoc* test. For the frequencies that ANOVA was significant in power and coherence, we performed a Holm-Sidak multiple comparisons correction. Statistical significance was set at p < 0.05.

## Results

### Demographics

Thirty-one dogs were evaluated for this study. Four were not enrolled because of a history of epileptic seizures, one dog was excluded for having elevated plasma levels of creatinine and one dog was excluded because it was euthanized a month later and necropsy revealed a brain tumor. Of the remaining 25 dogs, 12 were classified as normal with a median RDS of 9.5 (Range: 9 – 12), 7 as at risk of developing CCD with a median RDS of 17 (Range: 15 – 21) and 6 with CCD with a median RDS of 26 (Range: 23 – 33). Sex, age, and breed distribution for each group is shown in table 1. Dogs at risk and with CCD were significantly older than normal dogs (p = 0.0021 and p < 0.0001 respectively). No age differences were found between dogs at risk and dogs with CCD (p = 0.1438).

**Table 1.** Sex, age and breed distribution of dogs in each group according to their Rofina questionnaire score.

|  | Normal | At risk | CCD |
| --- | --- | --- | --- |
| **Sex** | 2 Intact males<br>4 Castrated males<br>6 Female spayed | 3 Intact males<br>1 Castrated male<br>3 Females spayed | 3 Intact males<br>3 Females spayed |
| **Age (years)** | 11.5 (8 – 14) | 13 (12 – 16) | 14.5 (11 – 17) |
| **Breeds** | 9 mix-breed<br>1 Labrador retriever<br>1 Border collie<br>1 Pitbull | 4 mix-breed<br>1 Border collie<br>1 Beagle<br>1 Schnauzer | 4 mix-breed<br>1 Boxer<br>1 toy poodle |

### Dogs at risk of CCD have higher alpha power at the frontopolar cortex

Figure 1B shows a raw EEG in all the left electrodes of one of the CCD dogs. We evaluated the differences in the power of each frequency band between the three groups of dogs (Figure 2). The ANOVA analysis showed significant differences between groups for the alpha frequency band at Fp1 (F_2,22_ = 4.04, p = 0.032). Dogs at risk had higher alpha power than normal dogs (adj p = 0.033) but not than dogs with CCD (adj p = 0.735). Additionally, dogs with CCD showed no significant differences from normal dogs (adj p = 0.212). No other differences were observed between groups at any frequency band in any electrode location (adj p > 0.05). As it can be observed in Figure 2, dogs with CCD showed a peak of power at sigma frequency band. However, the peak was mainly caused by one of the dogs and no significant differences were found between groups.

**Figure 2.**
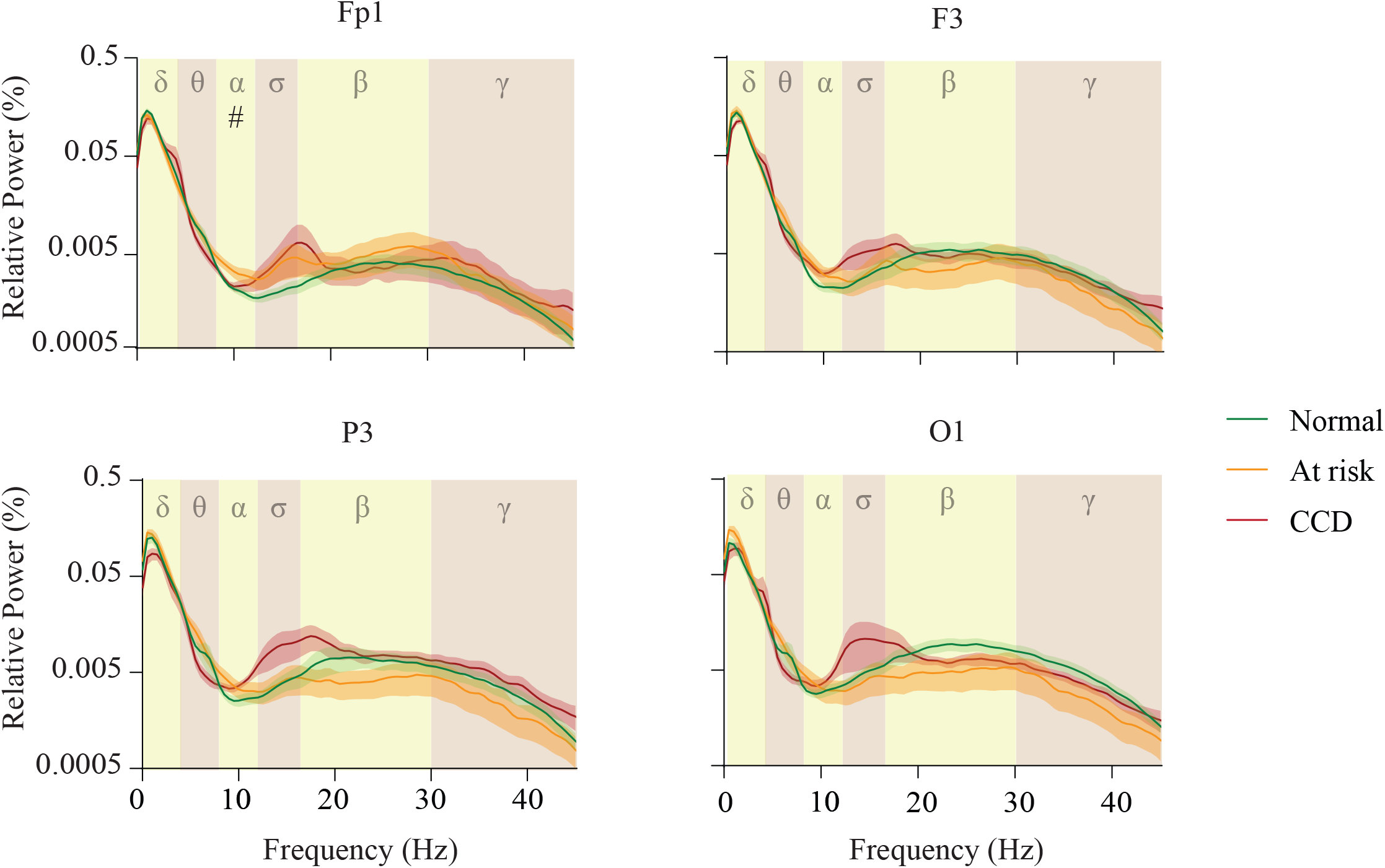
Power spectral profiles. Mean (solid line) and SEM (shadow) relative power spectral profiles of the left hemisphere in normal dogs, at risk, and with CCD. The analyzed frequency bands are indicated by different colors and their Greek letters in the background of the graphics. Significant differences are indicated by symbols: * Normal Vs. CCD, † Normal Vs. at risk, # At risk Vs. CCD.

### High frequency interhemispheric coherence is impaired in dogs with CCD

We determined the interhemispheric and intrahemispheric coherence between consecutive electrode locations (Figure 3A). The ANOVA analysis demonstrated differences between groups in the interhemispheric alpha coherence for P3-P4 (F_2,22_ = 6.616, p = 0.0056), beta coherence for F3-F4 and P3-P4 (F_2,22_ = 4.007, p = 0.0328 and F_2,22_ = 3.499, p =0.0479) and gamma coherence for Fp1-Fp2, F3-F4, P3-P4 (F_2,22_ = 3.980 p = 0.335, F_2,22_ = 5.091, p = 0.0152, F_2,22_ = 4.651, p = 0.0207 respectively). After multiple comparisons correction we demonstrated that dogs with CCD had lower gamma Fp1-Fp2 and F3-F4 coherence and lower beta F3-F4 coherence than normal dogs (adj p = 0.0347, adj p = 0.0255 respectively and, adj p = 0.0255), and dogs at risk had higher alpha P3-P4 coherence than dogs with CCD (adj p = 0.0363). Additionally, in the individual analysis, beta and gamma P3-P4 coherence were lower in CCD dogs than in controls (p = 0.017 and p = 0.0223), but the significance was lost with multiple comparisons correction (adj p > 0.05).

**Figure 3.**
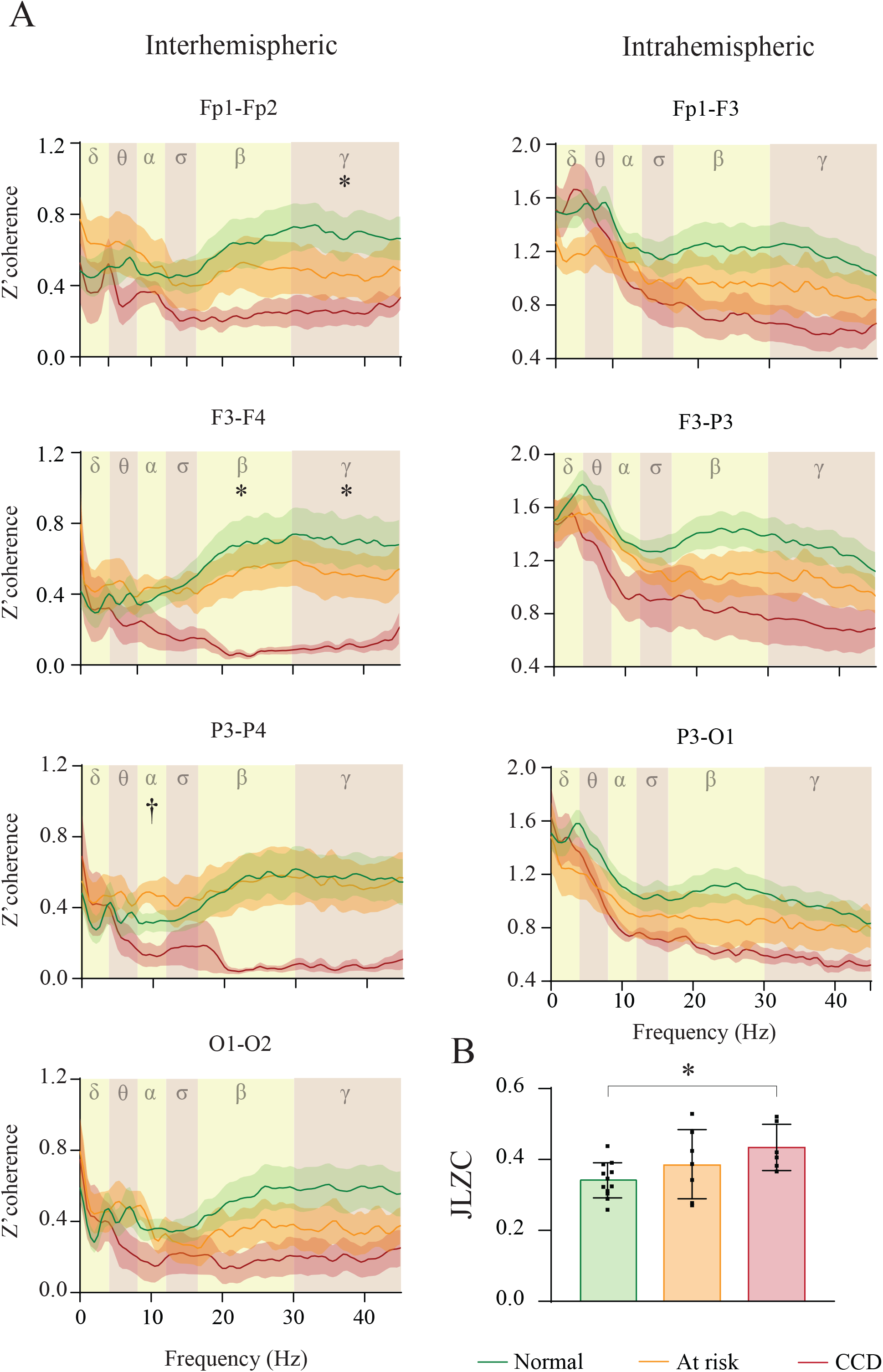
Z’ Coherence and Joint Lempel Ziv Complexity. **A.** Mean (solid line) and SEM (shadow) interhemispheric (left graphics) and intrahemispheric (right graphics) Z’coherence between adjacent electrodes in normal dogs, at risk, and with CCD. The analyzed frequency bands are indicated by different colors and their Greek letters in the background of the graphics. Significant differences are indicated by symbols: * Normal Vs. CCD, † Normal Vs. at risk, # At risk Vs. CCD. **B.** Comparison of Joint Lempel Ziv Complexity (JLZC) between the group of dogs. Asterisks represent significant differences.

### Increased temporo-spatial complexity characterizes dogs with CCD

We compared the LZC at each left electrode location, and we did not find any differences between groups (Table 2). However, when we analyzed the global complexity of the brain by means of JLZC (Figure 3B), we found groups to be significantly different (F_2,22_ = 3.689, p = 0.0415) with higher JLZC in dogs with CCD in comparison to normal dogs (adj p = 0.0356).

**Table 2.**
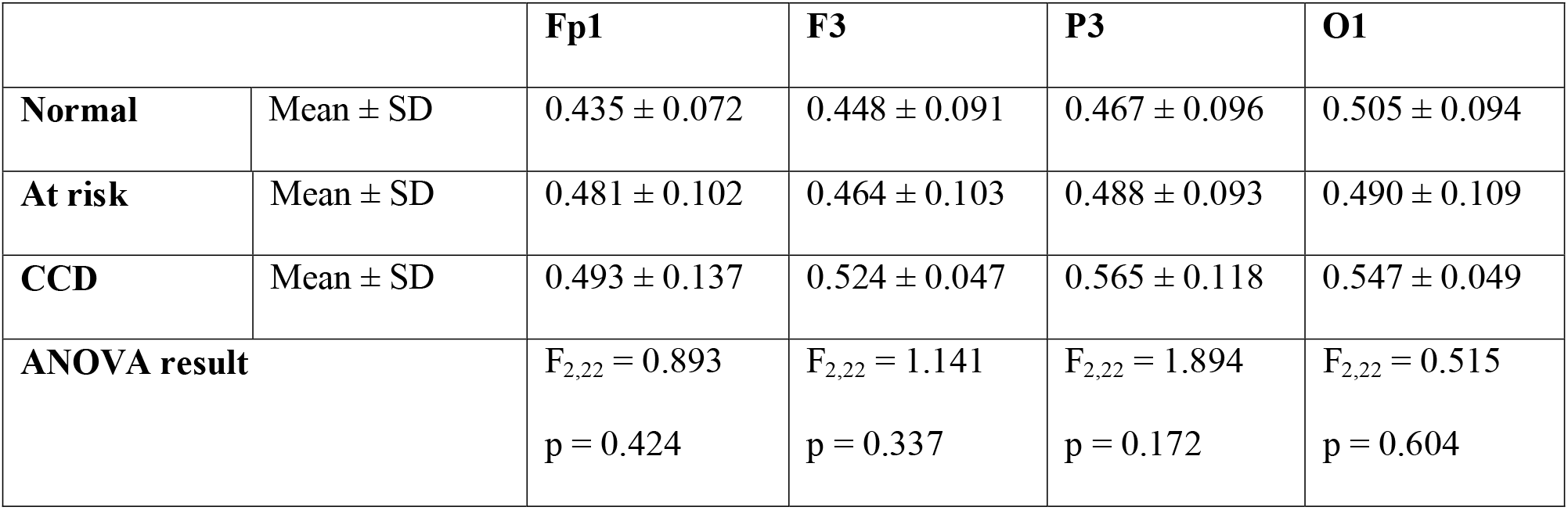
Lempel Ziv Complexity at each electrode location for each group of dogs. Descriptive statistics and one-way ANOVA results.

|  |  | **Fp1** | **F3** | **P3** | **O1** |
| --- | --- | --- | --- | --- | --- |
| **Normal** | Mean ± SD | 0.435 ± 0.072 | 0.448 ± 0.091 | 0.467 ± 0.096 | 0.505 ± 0.094 |
| **At risk** | Mean ± SD | 0.481 ± 0.102 | 0.464 ± 0.103 | 0.488 ± 0.093 | 0.490 ± 0.109 |
| **CCD** | Mean ± SD | 0.493 ± 0.137 | 0.524 ± 0.047 | 0.565 ± 0.118 | 0.547 ± 0.049 |
| **ANOVA result** | | $F_{2,22} = 0.893$<br>$p = 0.424$ | $F_{2,22} = 1.141$<br>$p = 0.337$ | $F_{2,22} = 1.894$<br>$p = 0.172$ | $F_{2,22} = 0.515$<br>$p = 0.604$ |

## Discussion

We have shown that in a manner similar to humans with AD, dogs with CCD and at risk of CDD have changes in the electrical activity of the brain. We demonstrated changes in EEG power, coherence and complexity.

### EEG power

Dogs at risk of CCD, but not dogs with CCD had higher levels of alpha power at the prefrontal cortex. This result suggests that in the early stages of the disease, compensatory mechanisms to maintain cognitive functions might take place. This has been also proposed in previous studies in humans. While most studies have found a decrease in alpha power in AD patients (Babiloni et al., 2009; Schreiter Gasser et al., 2008), Gauber et al. (2019) showed that in preclinical stages of AD, EEG patterns are modulated differently depending on the degree of amyloid deposit (Gaubert et al., 2019). They showed that the relationship between high frequency oscillations and amyloid burden followed an inverted U-shape, i.e., up to a certain level, higher levels of amyloid deposit correlate with higher frequency oscillations power, but when amyloid load exceed a particular threshold, the relationship inverts. Along the same lines, Nakamura et al. (2018) using magnetoencephalography showed increased frontal alpha power in early stages of AD (Nakamura et al., 2018). Similarly, in our study, dogs at risk of CCD exhibited higher alpha power only at the prefrontal cortex, which has been shown to typically be the site of disease onset in dogs (Prpar Mihevc and Majdic, 2019). According to our results, alpha power increases might be a useful marker of early stages of CCD.

Something to consider is that human studies are usually performed in resting eye-closed conditions and the alpha rhythm is characteristic of this state(Markand, 1990). In those studies subjects were asked to remain calm with eyes-closed during the recordings, however this is not feasible in non-trained dogs, and in this study, dogs remained with eyes open for most of the recording time. In humans, opening eyes leads to a significant reduction of alpha power caused by desynchronization of neuronal activity. This process is known as alpha reactivity, and it is considered a marker of cholinergic system integrity (Schumacher et al., 2020). Alpha reactivity is impaired in AD and other dementias such as Lewy Body; demented subjects show higher values of alpha power than controls (Babiloni et al., 2010; Schumacher et al., 2020). In dogs, an age-related deterioration of the cholinergic system functions that correlates with cognitive decline has been demonstrated (Araujo et al., 2005), therefore an impairment in alpha reactivity is expected and could also explain the observed alpha power increase in dogs at risk.

In contrast to humans with AD, dogs with cognitive dysfunction did not show a slowing of their EEG activity; i.e., an increase in delta and theta oscillations(Gianquinto and Nolfe, 1986; Moretti, 2004). However, the slowing of the EEG seen in AD occurs mainly in severely demented patients and preclinical stages can even show decreases of delta power (Gaubert et al., 2019). Further studies with a higher number of severely affected dogs are needed to evaluate whether slowing of lower frequencies can be observed at later stages of the disease.

### EEG coherence

We demonstrated a reduction in the interhemispheric coherence of beta and gamma oscillations at the frontopolar and frontal cortex in dogs with CCD. High frequency bands coherence plays an important role for information processing and cognition (Fries, 2015; Torterolo et al., 2019), and is severely impaired in patients with AD (Jelles et al., 2008; Koenig et al., 2005; Pogarell et al., 2005). The reduction in interhemispheric coherence in this study implies a loss of functional connectivity between both hemispheres. This coherence reduction is correlated with corpus callosum atrophy in humans with AD (Pogarell et al., 2005). This structure is the main bundle of nerve fibers that connect cerebral hemispheres. Age-related axonal degeneration within the corpus callosum have been demonstrated (Barry et al., 2021), but to the authors’ knowledge, differences in corpus callosum size between dogs with and without CCD have not yet been reported. Of note, we found a higher alpha coherence in dogs at risk than in CCD dogs. While values of dogs at risk were not significantly different from normal dogs, they were higher, which could suggest, again, that compensatory mechanisms are taking place in early stages of the disease.

### EEG complexity

While one of the characteristics of AD patients’ EEG is the reduction in complexity (Labate et al., 2013; Liu et al., 2016), we did not see group differences in LZC in any electrode location. However, when we analyzed the global complexity of the brain by means of JLZC, dogs with CCD had higher complexity than normal dogs. This result could be explained by the loss of synchronization between different cortical areas as shown with the reduction in coherence. Comparably, omega complexity (a measure of spatial complexity of the whole brain) is also higher in patients with AD in comparison with controls suggesting a desynchronization of different brain regions. In this sense, JLZC is a novel measure that provides a broader analysis of cortical complexity and appears to be more useful than complexity at each electrode site in diseases characterized by loss of homogeneity across cortical areas.

### Limitations and future directions

One of the biggest limitations in CCD research at this time is the lack of specific diagnostic tests other than autopsy. As such, client questionnaires have become the accepted means of recognizing the typical neurobehavioral signature of the disease, coupled with physical and neurological examinations as well as blood work to rule out other causes. However, it is important to note that the underlying pathology in the dogs reported here has not been confirmed. Ages were not perfectly matched between groups, i.e., dogs in CCD group were older than in the normal group. Therefore, some of the differences between the groups of dogs might not be due to their cognitive status only, but also to the effects of healthy aging. However, as prevalence of behavioral changes associated with cognitive dysfunction increases with age, reaching levels of 68% in dogs 15-16 years old (Neilson et al., 2001), finding cognitively healthy dogs in the higher limit of age becomes challenging. Additionally, EEGs were recorded during wakefulness and dogs kept their eyes mainly open, making hard the comparison with humans’ recordings during the eyes-closed resting state. Finally, we have used owners’ questionnaires, an indirect measure of dogs’ cognitive dysfunction, but we did not perform any direct measure such as cognitive testing.

Future studies should evaluate CCD EEG features during sleep, since NREM (Bonnani et al., 2012; Taillard et al., 2019) and REM (Hassainia et al., 1997; Petit et al., 2004) sleep correlates of AD have been also demonstrated, and some of them are more specific than the ones find during wakefulness (Petit et al., 2004). Interestingly, it would be desirable to employ a non-invasive polysomnography method, that has recently been described and validated (Gergely et al., 2020; Kis et al., 2014).

## Conclusions

We have demonstrated that dogs diagnosed presumptively with CCD show one of the main changes observed in AD patients, the loss of gamma coherence, suggesting a decrease in functional connectivity between hemispheres. Furthermore, CCD patients showed an increased JLZC. Additionally, dogs at risk of CCD had higher alpha Fp1 power and P3-P4 coherence which could be related to compensatory mechanisms that take place at the early stages of the disease. These results suggest that alpha power and coherence could be useful in the early diagnosis of CCD. In addition of the importance of these results for understanding this canine disease, our findings reinforce the domestic dog as a suitable translational model of AD. Longitudinal studies are needed to evaluate how changes in cognitive capacities correlate with shifts in quantitative EEG over time.

## Acknowledgements

The authors would like to thank all the owner’s that agreed to have their dogs participate in this study. We are grateful with Dr. Katharine Russell for her help with figures. This study was supported by Comisión de Investigación y Desarrollo Científico (CIDEC), Facultad de Veterinaria, Uruguay.

## Declarations of interest

None

## Notes

### Competing Interest Statement

The authors have declared no competing interest.

